# Convergent evolution in silico reveals shape and dynamic principles of directed locomotion

**DOI:** 10.1101/2022.11.20.516954

**Authors:** Renata B Biazzi, André Fujita, Daniel Y Takahashi

## Abstract

Active, directed locomotion on the ground is present in many phylogenetically distant species. Bilateral symmetry and modularity of the body are common traits often associated with improved directed locomotion. Nevertheless, both features result from natural selection, which is contingent (history-dependent) and multifactorial (several factors interact simultaneously). Based solely on the unique natural history on Earth, it is difficult to conclude that bilateral symmetry and modularity of the body are required traits for an improved locomotion ability as they can result from chance or be related to other body functions. As a way to avoid these caveats, we propose using physics-based simulations of 3D voxel-based soft robots evolved under different evolutionary scenarios to test the necessity of both traits for sustained and effective displacement on the ground. We found that an intermediate number of body modules (appendages) and high body symmetry are evolutionarily selected regardless of gravitational environments, robot sizes, and genotype encoding. Therefore, we conclude that both traits are strong candidates for universal principles related to improved directed locomotion.

## Introduction

Active and directed locomotion is a ubiquitous ability necessary for the survival of several species. Animals have different locomotion due to the complex body-environment relationships developed during their evolutionary history (***Alexander, 2006; Biewener and Patek, 2018***). The shape and dynamics principles behind animals’ locomotion ability entangle with all other body’s functions and needs for survival (*e.g*., the origin of the fins-to-limbs transition of tetrapod is probably the result of multiple and interconnected selection pressures (***Hecht et al., 1977; Panganiban et al., 1997; Dickson and Pierce, 2018; Molnar et al., 2021***)). Thus, it is not straightforward to identify in an animal’s shape what are the necessary basic conditions for improved autonomous locomotion, and what could have been different (***Alexander, 2006; Shubin et al., 2009***). We cannot ascertain which of the evolutionary outcomes we see today would be robustly replicated when replaying the “tape of life” on Earth on potentially another planet (***Powell and Mariscal, 2015***).

Bilateral symmetry and body segmentation are two fundamental body plan features shared by most modern animals. The evolutionary origin of these two features is complex. Both features are associated with widely diverse genetic regulatory networks that combine homologous and convergent characteristics (***Erwin and Davidson, 2002; De Robertis, 2008; Baguñà et al., 2008; Couso, 2009; Manuel, 2009; Zhang et al., 2014; Holló, 2015; Chen et al., 2019***). The evolutionary origin of bilateral symmetry is often attributed to a fitness gain in directed locomotion (***Holló and Novák, 2012***) - the ability to continuously sustain displacement in one direction. However, it can also be associated with an improvement in internal transport in the body (***Finnerty, 2005***). Moreover, symmetric structures might generally prevail simply by best satisfying the restrictions imposed by the physical laws acting in a body (***Holló, 2017***). Likewise, the correlation between mobility and metamerism (***Couso, 2009***) and the evolution of appendages associated with changes in locomotion strategies (***Panganiban et al., 1997; Shubin et al., 1997***) suggests that a body divided into parts is relevant to locomotion (***Alexander, 1982, 2006; Hildebrand, 1988; Biewener and Patek, 2018***). Nevertheless, it is also likely that evolutionary pressure for solutions that minimize descriptional complexity for genotype-phenotype maps bias the selection to symmetric and modular structures (***Johnston et al., 2022***).

Using the ground - on land or inside the water - to support and propel the body is one of the most typical forms of animal locomotion (***Alexander, 2006; Biewener and Patek, 2018***). Prominent examples of this type of locomotion are animals that live outside water without being able to fly and non-sessile crustaceans. Beyond them, ground locomotion exists in fishes that evolved limb-like adaptations like frogfish (Figure 1A), batfish, and mudskippers and even in unexpected cases such as octopuses walking (***Dickson and Pierce, 2018; Amodio et al., 2021***). Thus, locomotion on the ground is present in phylogenetically distant species (such as the maned wolf and frogfish in Figure 1A) and depends upon one of the essential features of Earth and other planets on which life might appear - the presence of a floor. To better understand the evolution of body characteristics associated with locomotion, here we investigate whether symmetry and modularity (defined as the division in connected body parts that move together with relative independence of the rest of the body) are features that an organism’s shape *need* to have for optimizing average speed on the ground, independently of the specific evolutionary history and non-locomotion related evolutionary pressures. Figure 1B shows a schematic representation of symmetry and modularity on the maned wolf and frogfish bodies.

**Figure 1.**
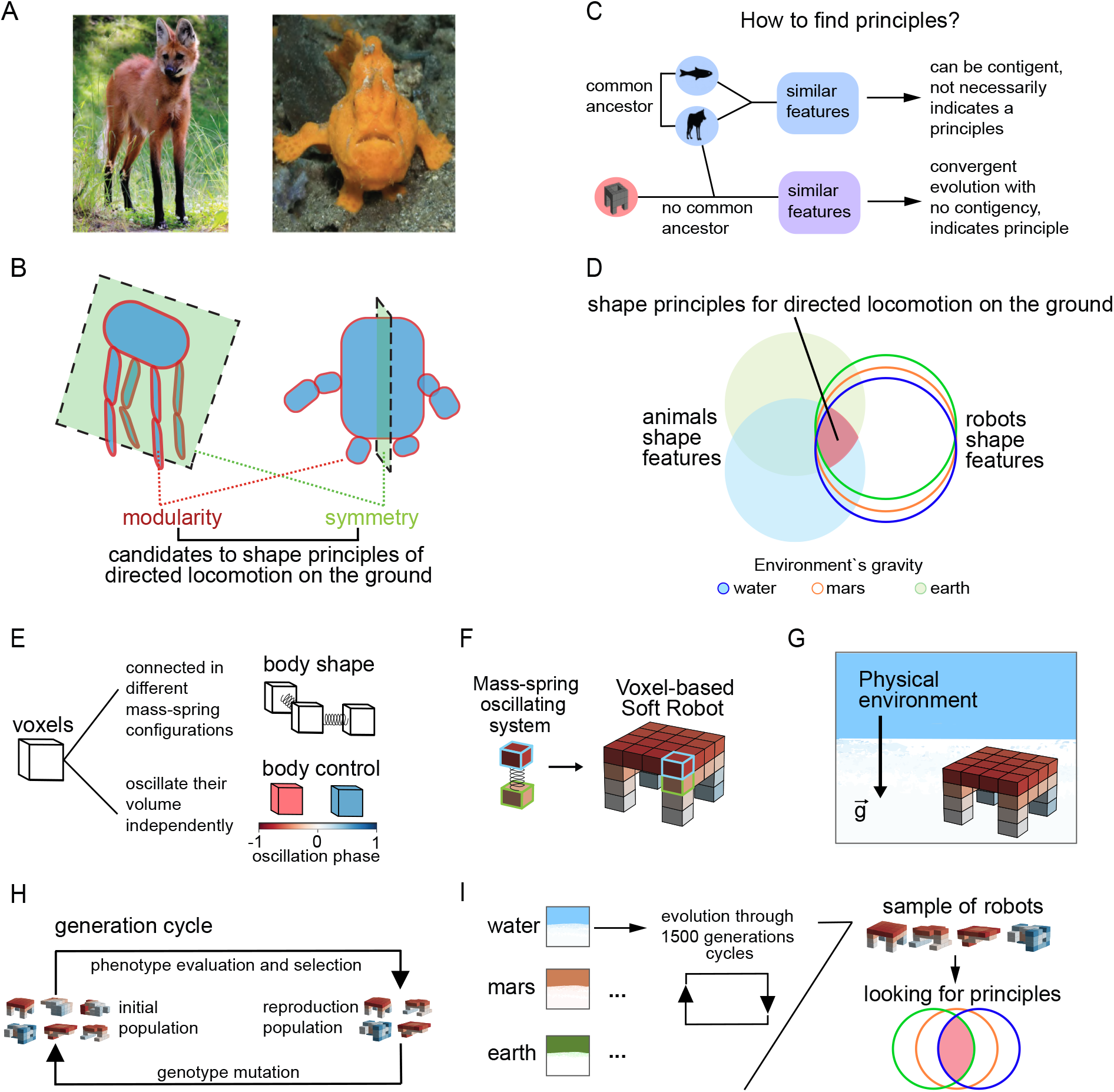
The convergent evolution of animals and robots makes it possible to find principles of locomotion without contingent dependence. (A) A maned wolf and a frogfish. Both species use the ground for locomotion in their environments. (B) Schematic illustration showing the symmetry and modularity of the maned wolf and frogfish bodies. The modules (blue outlined in red) are the joining connected body parts that move together during locomotion. The symmetry plane (green) indicates the bilateral symmetry of the body. Modularity and symmetry are present in phylogenetically distant species. Therefore, they are candidates for being necessary features (principles) of directed locomotion on the ground. (C) Convergent evolution of organisms with no common ancestor (*i.e*., animals and robots) and just one common behavior (*i.e*., locomotion) can be used to find principles of this behavior. It allows differentiating a necessary feature for enhanced locomotion from features resulting from the other animal’s functions or contingent on Earth’s evolutionary history. (D) The green and blue-filled circles represent the shape features of terrestrial and aquatic animals with directed locomotion on the ground. The green, orange, and blue unfilled circles represent, respectively, the shape features of robots with directed locomotion on the ground subject to 9.81*m/s*^2^ (Earth’s gravity), 3.721 (Mars’s gravity), and 0.1 (gravity plus buoyancy inside the water) acceleration towards the floor. The red intersection at the center represents the convergent features that indicate principles expected to be valid in different gravitational environments and organisms as different as animals and robots. (E) A voxel is the unit constituting the robots. The voxels are connected in different configurations (defining the body shape) and oscillate their volume independently (defining the body control), forming a mass-spring oscillating system. (F) Example of a Voxel-based Soft Robot constituted by mass-spring oscillating units. (G) We simulate the robots in a physical environment that evaluates effects like gravity, friction, and floor stiffness in the mass-spring system of voxels. (H) The generation cycle is the unit of the robots *in silico* evolutionary process. The phenotype evaluation is the average speed of the robot in its environment calculated in the 30s of simulation (directed locomotion ability). The best robots are selected, and their genotype (CPPN’s networks) are mutated to produce a new generation of robots that will constitute the next initial population. (I) In each environment (water, mars, and earth), an evolutionary process of 1500 generation cycles results in a sample of robots with different bodies (shape and control) and performances. The shape and control features of the robots are analyzed looking for principles. Photo credit: Two Fish Divers (frogfish) and Wikipedia user Calle Eklund/V-wolf (Maned wolf) (CC BY-SA 3.0).

There are two requirements to address how necessary symmetry and modularity are for the active and directed locomotion of animals on the ground. First, we need to separate locomotion from other animal needs and functions. Second, we have to access locomotion solutions that are not contingent on having N=1 evolutionary history. We propose that simulating an evolutionary process using robots under realistic physical constraints is a solution to satisfy the above requirements (Figure 1C). Robots allow us to test hypotheses about animals and search for principles of locomotion (***Aguilar et al., 2016; Gomez-Marin and Zhang, 2022***). In robotic simulations we can choose *which* biological process and physical constraints will be emulated, a controlled uncoupling of animal’s complexity that is usually difficult to obtain using other methods (***Ijspeert, 2001; Webb, 2002, 2009; Ijspeert, 2014; Karakasiliotis et al., 2016; Fukuhara et al., 2020; Schwab et al., 2021; Wenguang et al., 2021; Gomez-Marin and Zhang, 2022***). Also, in robot simulation we can have N>1 evolutionary histories, a necessary condition to study the contingency level of the outcomes. One crucial environmental parameter we can control in the simulations is gravity. It allows us to study the effects of different body gravitational loads in locomotion on the ground (***Rayner, 2003***). This way, we can address the question if external symmetry and modularity are robust principles of locomotion valid on the ground either inside the water (0.1*m/s*^2^ - almost no body load caused by buoyancy), on Earth’s surface (9.81*m/s*^2^), or in an imaginary martian life form (3.72*m/s*^2^). The convergent evolution of shape traits evolved independently in very different systems that have just the directed locomotion ability in common (animals and robots) would justify calling them principles of directed locomotion on the ground (Figure 1D).

## Results

For the simulation of an evolutionary process using robots, we choose to use 3D Voxel-based soft robots as our organisms (Figure 1F). We used the simulation engine proposed on ***Hiller and Lipson*** (***2014***) (Voxelyze). This software efficiently and accurately simulates soft bodies’ interaction with an environment based on physics laws. The body unit is the voxel. Voxels have features like mass, density, and stiffness and connect with other voxels as mass-spring systems forming the body shape (Figure 1E). The volume of each voxel oscillates cyclically and in an independent moment given by its oscillation phase (Figure 1E). The connected voxels work as muscle units as they expand, contract, and deform, and we can control their stiffness and density. The robots’ voxels were connected inside a space dimension of 4 × 4 × 4 possible voxel positions (see Material and Methods). We choose to first explore exhaustively the 4^3^ space dimension, as it is the minimal possible space that allows meaningful body plans. We also did control experiments within 6^3^ and 8^3^ to check for dimension size effects. We simulated the robots in a physical environment (Figure 1G) that models effects like gravity, floor stiffness, and friction to properly simulate collisions and damping between voxels and the environment (***Hiller and Lipson, 2014***). The design and locomotion behavior of evolved robots using Voxelyze successfully transferred to organisms made of biological tissue (***Kriegman et al., 2020***). We based our simulation parameters on these previous works.

We generate the robot’s sample using an artificial evolutionary process that selects for better locomotion ability - defined as higher average speed as it is a proxy for organisms with sustained and effective displacement (see Materials and Methods). This process contains the fundamental elements of natural evolution: i.reproduction, ii.passage of information through generations (hereditary inheritance), iii.phenotype variation by mutation in genotype and iv.selection of the fittest (Figure 1H). Three independent evolutionary processes of 1500 generations were simulated, one for each gravitational environment - 0.1 (water), 3.72 (mars), and 9.81*m/s*^2^ (earth). In the water-like environment, the bodies have nullifying body weight but do not have drag effects. We did not add drag in our simulations because our aim is to study just the body weight influences in locomotion independently of other forces. Each evolutionary process is a search in the space of shape and control solutions. After 1500 generations, we had a sample of robots with different bodies and directed locomotion abilities (average speed). This process was repeated 30 times (seeds) for each environment, and in the end, we had big samples of different robots for each of the three gravitational environments. In each environment, we classified every robot inside one of the five fitness layers (100%-80%, 80%-60%, 60%-40%, 40%-20% or 20%-0%) according to its fitness ratio relative to the best robot in that environment (100% fitness). With this sample, we tested the hypotheses about the relationships between locomotion performance and body modularity and symmetry (Figure 1I).

### An intermediate number of modules sparsely connected are necessary to increase directed locomotion on the ground

To study body modularity, we clustered the robot’s voxels based on their proximity and synchronization. We considered voxels that are connected and have a similar phase offset as being a member of the same module (cluster) (Figure 2A). A module has the average position and phase of its voxels members. The modules are linked if they have voxels directly connected. A topological representation - consisting of the robot’s modules and connections - is constructed for each robot (Figure 2A). We visualize the topological representation in two ways. The first one (TP1) colors the modules depending on their phase value and places them using their mean positions. The size of the module depends on the number of voxels belonging to it. The modules that touch the ground are represented as triangles, while those that do not are circles (see Figure Figure 2A). The second type of visualization (TP2) shows the modules and their connections without adding any other information (Figure 2A). The average degree measures how connected a network is. Figure 2B exemplifies different robots’ modularity and average degree value.

**Figure 2.**
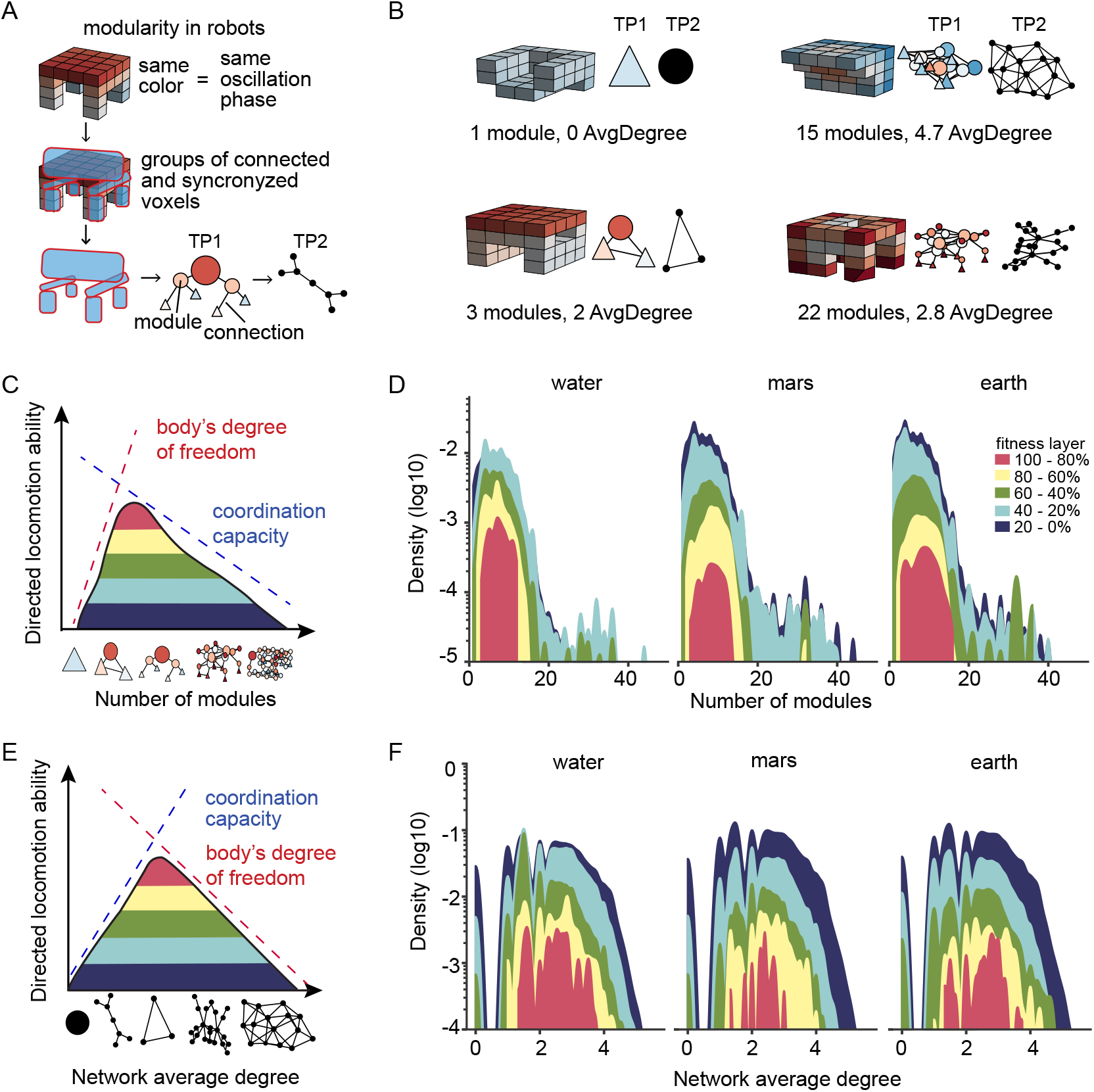
An intermediate number of modules sparsely connected is necessary for achieving higher directed locomotion performance. (A) The modules are groups of connected and synchronized voxels of a robot. Using the modules, we created two topological representations of the robot (TP1 and TP2). TP1 has information about the mean phase (represented by the color) and the position of each module. The size of each module is relative to the number of voxels it contains. Modules that touch the ground are represented as triangles, while those that do not are circles. TP2 representation shows the modules and their connections without adding any further information. (B) Robot’s modularity examples with the number of modules and the connectivity measure (AvgDegree). (C) Illustration of the hypothesized impact of the number of modules on the directed locomotion ability. The robots with the 20% top directed locomotion ability (red area) are the ones that maximize both the body’s degree of freedom and its coordination capacity by having an intermediate number of modules. (D) The results of the three environments confirm the hypothesis: the best robots (100-80% fitness layer in red) typically have an intermediate number of modules (more than one but less than 17). The module number range increases as the fitness layers worsen, representing the broad search. (E) Illustration of the hypothesized impact of the module’s connectivity (measured by the network average degree defined in Equation 1) on the directed locomotion ability. The robots with the 20% top directed locomotion ability (red area) maximize both the body’s degree of freedom and coordination capacity by having sparsely connected modules. (F) The results of the three environments confirm the hypothesis that robots with 20% better directed locomotion ability (100-80% fitness layer) typically have an average degree between 1.2 and 4 (sparsely connected). Robots with modules densely connected (more than four connections per module on average) do not have so good performances. **Figure 2–Figure supplement 1**. Results of the number of modules and average degree in experiments using other robot sizes and genotype. **Figure 2–Figure supplement 2**. Consistency test of the results of the number of modules. **Figure 2–Figure supplement 3**. Alternative limbs configuration for bilateral quadrupeds.

We hypothesized that the robot’s locomotion ability should be higher when it maximizes both body’s degree of freedom and its coordination capacity (Figure 2C). The number of possible movements of the body increases with the number of modules (red dotted line in Figure 2C). However, more movement possibilities will not necessarily add to a better locomotion performance. Increasing the number of modules reduces the chance of finding suitable coordination for locomotion (Figure 2C blue dotted line) (***Aoi et al., 2016***). Thus the robots with the top fitness value (red) are expected to be in the region of an intermediate number of modules to maximize both body’s degree of freedom and its coordination capacity. Supporting our hypothesis, Figure 2D shows that the robots that belong to the top 20% fittest ones (100-80% fitness layer) have exclusively a small to an intermediate number of modules (from 3 to 17, approximately), indicating low tolerance of the best robots to have one or two modules and to increase their number of modules indefinitely. On the other hand, as the fitness layer worsens, they often include robots with a lower and higher number of modules; observe that the worst fitness layers (40-20% in light blue and 20-0% in dark blue) have very similar shapes accounting for a broad range of number of modules. This pattern repeats in all three environments (water, mars, and earth), indicating robustness for different gravities.

Additionally to the number of modules, the robot’s body dynamics depend on the module’s connectivity. Here, we used the network average degree, defined in Equation 1, to measure the body’s connectivity. Modules more connected make a more tied structure, implicating a decrease in the body’s degree of freedom. Furthermore, as the modules’ movement affects their neighbors, a structure that is strongly tied will have higher coordination in its movement (Figure 2E). Thus, we hypothesized that the robots with the top performance (red area in Figure 2E) would maximize both body’s degree of freedom and its coordination capacity by having an intermediate degree of connectivity (*i.e*. a sparsely connected body). Figure 2F shows that in the three environments, the 20% fittest robots (100-80% fitness layer) have, as expected, an intermediate average degree between 1.5 and 4. On the other hand, the worst fitness layers more often include robots with a lower and higher average degree.

Figure 2D and Figure 2F show the results of robots constructed inside a 4^3^ space dimension and encoded by Compositional pattern-producing networks (CPPNs) (***Stanley, 2007***). CPPNs are a popular genotyping strategy that facilitates the generation of bodies structures with continuities and regularities (***Clune and Lipson, 2011; Cheney et al., 2013; Kimura et al., 2016; Corucci et al., 2018; Kriegman et al., 2020***). If the hypothesis of balance between the body’s degree of freedom and coordination is correct, we expect that the same pattern of an intermediate number of modules and average degree should hold for different robots’ genotypes and sizes. Testing other genotype strategies is necessary to verify if our findings are due to the locomotion selective pressure or are a byproduct of a specific genotype encoding. Thus, we repeated the experiments done with CPPNs using a Direct Encode genotype structure - an encoding strategy where the presence or absence of a voxel is independently chosen from each other, resulting in less bias. We also tested the effect of different dimension sizes evolving robots of 6^3^ and 8^3^ dimensions using CPPNs as genotypes. We found that in all these other experiments, a higher fitness also requires an intermediate number of modules and average degree (Figure 2–Figure supplement 1), meaning that the locomotion selection pressure leads to the same shape and dynamic principles of best performance regarding the exact encoding structure.

In all the experiments, the number of modules results have a density decay in the extremities of the distributions. We checked if the amplitude difference between the layers was significant and not due to extremely rare outliers present in the worst layers because of their bigger sample sizes. By a bootstrapping analysis controlling the sample size, we found that the best robots (100-80%) have a smaller amplitude of the number of modules when compared to the worst robots (Figure 2–Figure supplement 2).

### Highly symmetric body shape and a break of total symmetry in control are necessary conditions to increase directed locomotion on the ground

To quantify the robot’s symmetry, we defined two separate symmetry measures for its shape and control (Figure 1E). The shape symmetry is the percentage of symmetric voxels after mirroring the robot in the x and y-axis - see an example of shape symmetric voxels in Figure 3A. The XY shape symmetry value is the mean of these two values, in other words, the percentage of voxels that are x-y symmetric in their positions. To calculate the control symmetry, we considered the symmetry of the voxel’s position and phase Figure 3A. Thus, the XY control symmetry value is the percentage of x-y symmetric voxels in their positions and have equal oscillation phases. Figure 3B shows two examples of robots with their shape and control symmetries calculated for the x-axis (X), y-axis (Y), and their mean value (XY).

**Figure 3.**
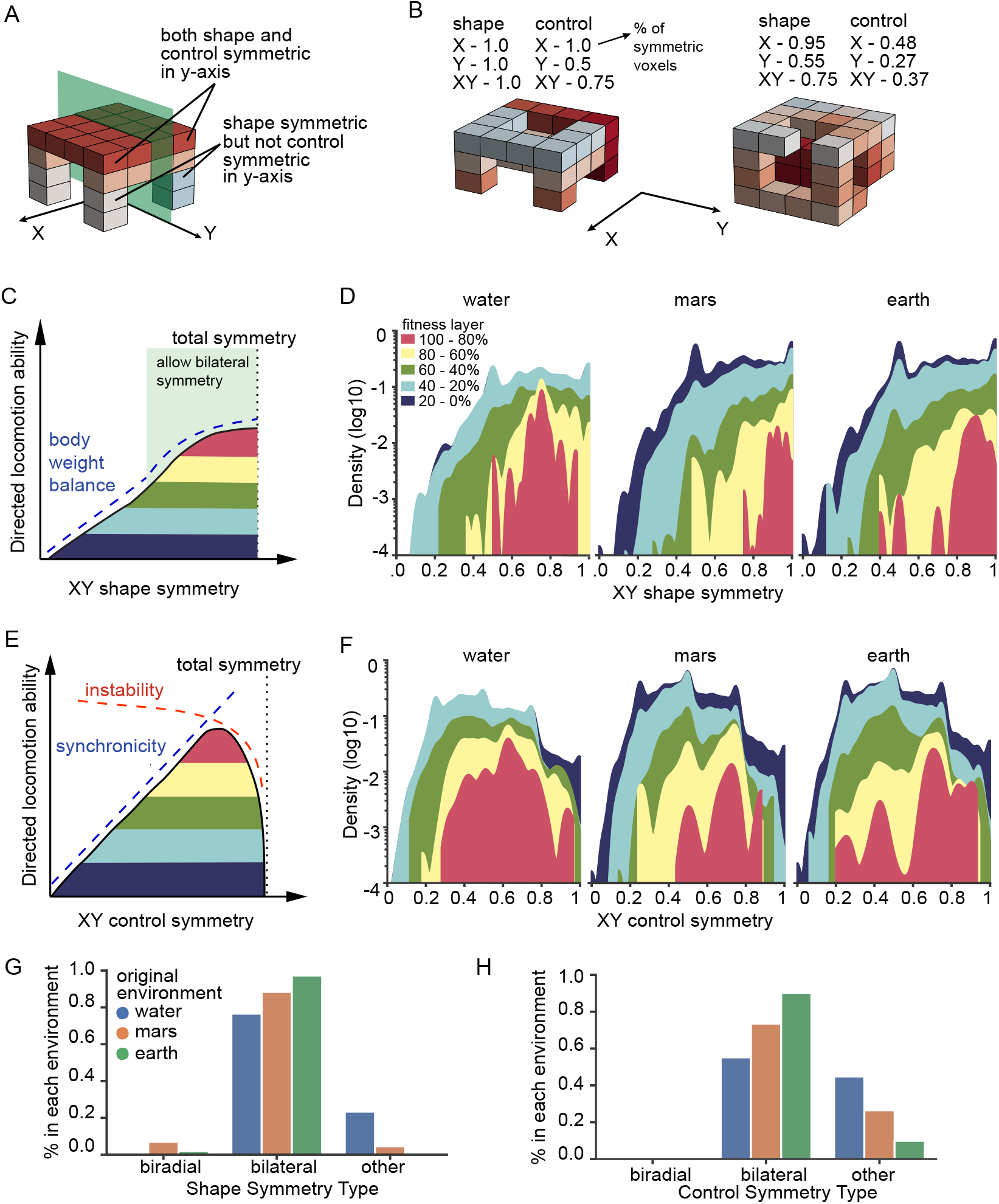
A highly symmetric shape and symmetry breaking in the dynamics are necessary for achieving higher displacements. (A) Illustration example of voxels with shape and control symmetry in the y-axis. (B) Examples of robot shape and control symmetry measures in the x-axis (X), y-axis (Y), and their mean value (XY). (C) Illustration of the hypothesized impact of XY shape symmetry on the directed locomotion ability. A poor body weight balance increases the body’s chance of falling or making a curved path. The green area allows bilateral symmetry in the body (100% symmetry at least on one axis). (D) In the results of the three environments, the robots with 20% better directed locomotion ability (100-80% fitness layer in red) typically have a shape symmetry higher than 0.5. The lower fitness layer contains a higher range of symmetry values. (E) Illustration of the hypothesized impact of XY control symmetry on the directed locomotion ability. The body synchronicity between its parts is higher with increased XY control symmetry. However, locomotion depends upon instability (break of symmetry) in the direction of the movement. Thus, the robots with top 20% performance (red) maximize synchronicity but necessarily keep a break of total control symmetry in the body. (F) In the results of the three environments, the robots with 20% better performance (red) have a peak shifted to higher values of symmetry and a break of total control symmetry (XY control symmetry<1). (G) The bilateral symmetry is the type of shape symmetry most frequent in the top robots (100-80% fitness layer). Biradial symmetry is also present. (G) The bilateral symmetry is also the most frequent type of control symmetry in the top robots (100-80% fitness layer). Biradial control symmetry is not present in this layer. Figure 3–Figure supplement 1. Symmetry results of experiments using other robot’s sizes and genotype.

There are three main types of locomotion sequences: *i*. tumbled body and no significant trajectory (stands in the same place), *ii*. tilted body and curved trajectory, and *iii*. stable body and straight trajectory. The symmetry of the shape impacts the probability of having one of these locomotion sequences by affecting the distribution of body weight. As the robots do not have sensory feedback abilities, the weight balance is defined as the body’s movement due to gravity forces (consequences of the weight distribution and surface contact points) (***Benda et al., 1994***). We hypothesized that the robots with the best directed locomotion ability would tend to have a symmetric body shape. A robot with a low XY shape symmetry (XY shape symmetry < 0.5) has a higher chance of having a poor weight balance, increasing the chance of the body tipping over, thus leading it to a lousy locomotion performance (blue dotted line in Figure 3C). As XY shape symmetry increases (XY shape symmetry > 0.5), the locomotion performance increases and starts to saturate when the XY shape symmetry approaches 1. A body does not need to be 100% symmetry in the x and y-axis to have a good balance for locomotion. Values of XY shape symmetry larger than 0.5 allow bilateral symmetry in the body (green area in Figure 3C). Supporting our hypothesis, the results show that in the three environments, the robots with top 20% fitness have their distribution shifted to higher values of XY shape symmetry (red areas in Figure 3D). The region of low shape symmetry (XY shape symmetry < 0.5) contains mostly robots from the smaller fitness layers (40-20% in light blue and 20-0% in dark blue areas in Figure 3D). Thus, bodies with low shape symmetry cannot acquire a locomotion performance as good as the ones with high shape symmetry.

Beyond the shape symmetry, the body’s dynamic symmetry (measured by the XY control symmetry) will also impact the locomotion ability. We hypothesized that increasing the XY control symmetry impacts positively on fitness as it describes the body synchronicity of movement (blue dotted line in Figure 3E). Nonetheless, locomotion requires a minimum instability - the dynamic possibility of translating the center of mass - in the direction axis to generate the necessary forward displacement (***Bruijn et al., 2013; Nagarkar et al., 2021***). The instability tends to be higher the more asymmetric the XY control (red dotted line in Figure 3E). Thus, we expect a break of total XY control symmetry to be a necessary condition for the robots with the top best 20% fitness (Figure 3E). Supporting this hypothesis, our results show that in the three environments, the distributions of the best robots (red in Figure 3F) have their peaks shifted to higher symmetry values (XY control symmetry > 0.6) but do not reach total symmetry (XY control symmetry = 1). In contrast, the 20-0% (dark blue) and 40-20% (light blue) fitness layers distributions occupy most of the [0,1] interval of possible XY control symmetry values. Thus, we see the requirement of XY control symmetry < 1 for optimizing locomotion.

Animals that use directed locomotion on the ground often exhibit bilateral symmetry, which biases the locomotion to a specific direction (in contrast to radial symmetry) (***Holló, 2015***). We asked if the top robots have a preferred shape direction (bilateral symmetry) in addition to their high shape symmetry. We classified as bilaterally symmetric the shapes and controls that are 100% symmetric in only one of the axis (X or Y) and as biradially symmetric the ones that are 100% symmetric both on the X and Y axis. The bodies that did not fit into either of these two categories belong to the “other” group. Figure 3G shows the shape symmetry classification and Figure 3H the control symmetry classification of the robots from the 100-80% fitness layer. Bilateral symmetry is the most common type of shape and control symmetry between the robots with the best locomotion performance. Biradial symmetry, a symmetry found in corals (sessile animals) and hydras (***Savriama and Klingenberg, 2011; Watanabe et al., 2014***), is a kind of shape symmetry present in a small portion of the top robots (Figure 3G).

As in the modularity case, we tested if our shape and control symmetry hypotheses are also robust to the other genotypes and robot sizes to ensure the results are due to the selective pressure and not a byproduct of specific scales or encoding choices. Figure 3–Figure supplement 1 shows that in all the experiments the best robots are shifted and restricted to higher shape and control symmetry values when compared to the other layers.

### Shapes are specialized to their gravitational environment

The best robots of the three environments have similar modular and symmetry features (Figure 2 and Figure 3) - thus justifying to call these features gravity-invariant principles. Nevertheless, bodies submitted to different gravities might still require distinct features optimized for their locomotion. To test this, we verified if a body shape optimized to one gravity value will keep its performance in another gravity value (Figure 4A). We selected the best 50 shapes (just voxel arrangement) of each environment and transferred them to the other two environments (see Material and Methods). In the new environment, the robots could test different control patterns for their fixed shapes until finding the dynamics that maximize their displacement (Figure 4B). One possibility is that the shapes have specializations selected to work well in their original environments but not in the others. In this case, we expect that the transference quality will decrease with the increase of gravity difference between the original and the new environment (Figure 4C). Figure 4D shows that the robots originally from water (blue) when in other environments do not have a performance as good as the robots originally evolved in mars (orange) or earth (green). Similarly, the robots originally from mars and earth do not perform as well in water as those that initially evolved in water.

**Figure 4.**
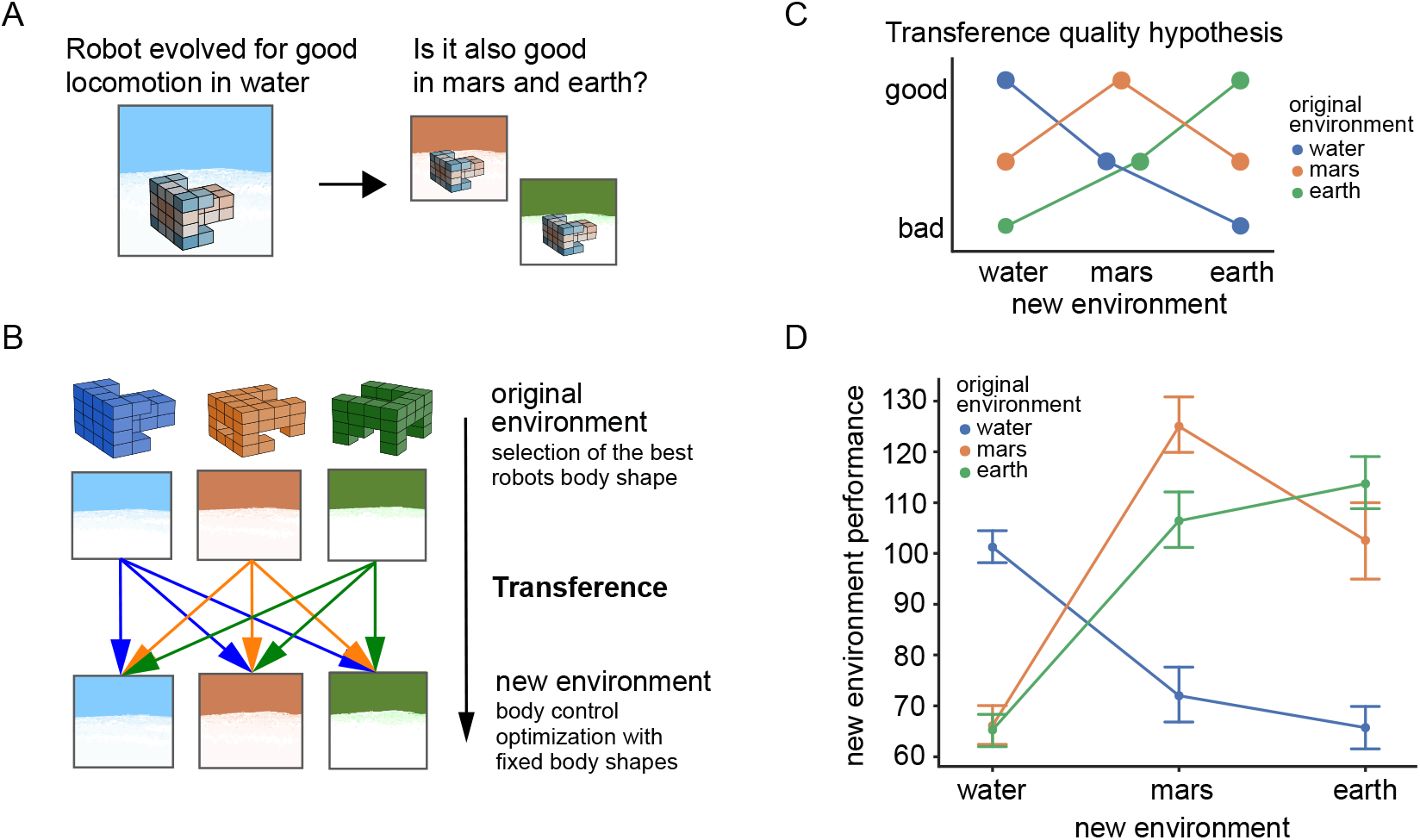
Robots from different gravitational environments require distinct features for good locomotion performance. (A) Will the best robots in water be the best when transferred to mars or earth? (B) Transference protocol. We optimize the best 50 body shapes (just shapes without control) of each original environment in the new environment. In each new gravitational environment, the fixed shapes could test different controls of their body shape and keep the best. (C) Illustration of a hypothetical transference outcome in which the quality of transference (the ability of directed locomotion in the new environment) will depend on the gravitational difference between the new and original environments. (D) The transferred robots’ average new environment performance (directed locomotion ability). Robots originally from water (blue) cannot acquire a performance as good as the best robots originally from mars (orange) and earth (green) in the mars and earth new environment. Robots originally from mars and earth have worse locomotion when transferred to water and cannot move as well as a robot originally from water.

#### Different gravity select different body structures

The shape’s lack of transference to other gravitational environments is due to the robot’s shape traits specialization for their original environments. A natural question is whether the robots in each environment have common traits. For example, one crucial component of the robot’s locomotion is the modules that touch the floor (Figure 5A). We posit that the relative size of the modules that touch the floor compared to the rest of the body has a different effect depending on the gravity. We found that the robot’s feet proportion distribution is different for each environment (*ANOVA test with p < 0.01, Figure 5B). The robots that originated from a lower gravity tend to have proportionally heavier feet (modules that touch the ground) than robots that originated from higher gravities (Figure 5B). To check if the feet’s proportion is related to the lack of transference between environments, we analyzed if there is a correlation between each robot’s feet’s proportion and its transference capability. We measured the transference capability as the difference in performance between the new and original environment. There is a negative correlation between the proportion of feet voxels and the robot’s locomotion transference capability when the robots go to an environment with higher gravity, *i.e*., water to mars (dark blue in Figure 5C), water to earth (light blue), and mars to earth (red) - with Spearman correlation coefficients of r = -0.39, r = -0.43, and r = -0.32 respectively, all with p < 1e-08. This result implies that lighter feet are usually better in higher gravity environments when compared to heavier feet.

**Figure 5.**
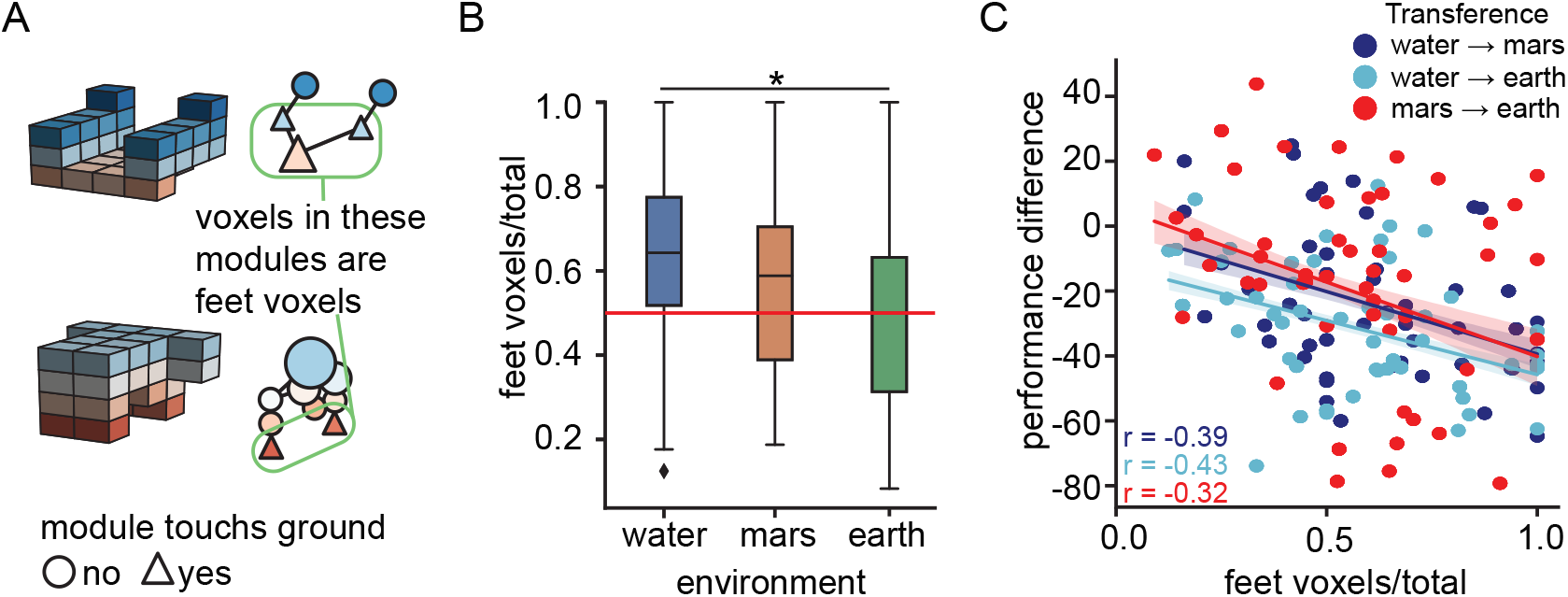
The relative volume of the feet affects robot performance when it goes to higher gravitational environments. (A) The voxels belonging to modules touching the ground during the robot’s movement are called *feet voxels*. (B) The distribution of feet voxels proportion in the body differs between the environments (*p < 0.01, ANOVA) and tends to smaller values when gravity increases. (C) In the environmental transitions that gravity increases - water to mars (dark blue), water to earth (light blue), and mars to earth (red) – the robots with a small proportion of feet voxels maintain their performance better (smaller difference between the new and the original environment). Spearman correlation coefficients of r = -0.39 (water to mars), r = -0.43 (water to earth), and r = -0.32 (mars to earth), all with p < 1e-08.

## Discussion

Directed locomotion on the ground is present in phylogenetically distant species. We used evolved robots to study which animal’s shape features are necessary to maximize this ability and which could be different without disrupting locomotion capacity. We found that the number of modules should be small and sparsely connected. The body’s shape should be highly symmetric, and the body control should exhibit symmetry breaking. Bilateral symmetry was shown to be the most successful type of symmetry. We also found that different gravitational environments require different shape structures to optimize locomotion average speed. Finally, we found that the feet’s proportion in the body is related to how well the bodies adapt to a higher gravitational environment.

### Morphological computation principles of directed locomotion on the ground

Biological systems carry out computations with their physical bodies to successfully interact with their environments - a concept called morphological computation (***Chiel and Beer, 1997; Tytell et al., 2011; Hauser et al., 2014; Zhang and Ghazanfar, 2018***). In this work, we investigated the existence of morphological computation principles relevant to directed locomotion on the ground. We used a population analysis of different synthetic organisms not restricted to the known biological solutions.

Animals have morphological modularity - a segmentation of the body related to its functional roles (***Williams and Nagy, 2001; Klingenberg, 2008; Larouche et al., 2018***). The number of modules, their size, and their positions are major features defining the types of locomotion of an animal (***Neil H***. ***Shubin and Marcus C***. ***Davis, 2004***). In our work, the modules are the connected body parts with little or no internal relative motion. The robot’s number of modules represents the number of different dynamic parts of an animal’s body - the trunk structure and its number of appendages and limb segments. We found that the optimization of locomotion necessarily requires a small to intermediate number of modules. A three-segmented limb structure, for example, is already sufficient to allow an organism to displace itself (***Ritzmann et al., 2004***). Indefinitely increasing the number of modules will saturate and possibly worsen their contribution to locomotion (***Aoi et al., 2016***). The modules also need to be sparsely connected, a type of connectivity that generates extremities. This way, our results are consistent with what we observe in different animal species with directed locomotion capacity - the presence of appendages (extremities) and a relatively small number of limbs and limb segments - both usually smaller than ten (***Fischer and Blickhan, 2006; Bruce, 2021***). This consistency is evidence that a small number of sparsely connected modules is a morphological computation principle for an organism’s optimized average speed.

Our work shows that high shape symmetry and a control (dynamic actuation) with a break of total symmetry is necessary for optimizing directed locomotion performance in an organism (***Nagarkar et al., 2021***). Specifically, bilateral symmetry was the most frequent type of symmetry among our top best robots. In animals, the bilateral symmetry type is present in more than 99% of the species (***Manuel, 2009; Holló, 2015***). An argument for the presence of bilateral symmetry is its equal favoring of both sides of the body in rectilinear motion in water (where life originated in the presence of drag forces) and its facilitating action in maneuverability by rapid changes of direction (***Holló and Novák, 2012***). Our results point out that shapes with bilateral symmetry are the best solution, even without historicity, maneuverability, and the presence of drag forces. This type of symmetry establishes a preferential direction in the organism’s body, which is a morphological computation contribution to directed locomotion. Furthermore, locomotion requires a dynamic asymmetry (instability) in the direction of movement (***Holmes et al., 2006; Aoi et al., 2016; Nagarkar et al., 2021***). A biradial symmetric shape achieves this instability only if it has an asymmetrical control pattern in the axis of movement. Meanwhile, a bilateral symmetric shape acquires instability by its shape asymmetry in the displacement axis. Thus, bilateral symmetry also simplifies and even enlarges the control effects on the body’s locomotion by its asymmetry in the direction of movement.

We found morphological principles that are robust to different gravitational environments. However, an organism subject to different gravitational loads still needs to modify its limbs coordination to have efficient locomotion (***Gillis and Blob, 2001; Minetti, 2001***). Beyond requiring different types of gait, gravity is an evolutionary force that influences the shape outcomes during an evolutionary process (***Rayner, 2003; Miras and Eiben, 2019***). We found that the robots have shape specializations to their gravitational environments as they cannot equally transfer to other environments, even being able to modify their movement coordination. Specifically, we found that a smaller feet’s proportion in the body correlates with better adaptability in higher gravitational environments. A possible explanation for this effect is that robots with a smaller feet’s proportion operate similarly to a Spring-Loaded Inverted Pendulum (SLIP) template (***Blickhan and Full, 1993; Holmes et al., 2006***). In the SLIP dynamics, a small foot connects to a longer leg with the body’s center of mass (CM) at its end, thus reducing friction effects and amplifying the CM horizontal displacement as it is farthest from the ground. On the other hand, robots with proportionally heavier feet can have higher body friction (more surface contact with the ground) and a CM closer to the ground. This way, they will have other types of dynamic propelling, like crawling, which is less suitable for higher gravitational loads.

All the results presented here were robust to different gravitational environments, genotype encoding, and phenotype size - even so, more tests in these aspects would still benefit the robustness of our results. Beyond that, extending the tests for other important aspects of locomotion behavior - such as noise on the ground, energetic costs, and maneuverability - by using other locomotion metrics - as energy efficiency, stability margin, and dissipated power (***Paez and Melo, 2014; Aoi et al., 2016***) - would also be relevant to evaluate the principle’s robustness.

### Contingency of evolutionary outcomes

An animal’s body has functions and needs beyond locomotion for its survival. The body results from an evolutionary process, defined by historically dependent selection factors and genetic heritage. In general, we cannot separate these factors. Here we investigate how a specific functional cause - optimization of average speed during directed locomotion on the ground - externally defines the phenotypic space of shape possibilities. This approach allows evaluating the contingency level of shape outcomes (***Powell and Mariscal, 2015***), impacting our expectations about the robustness of Earth’s animal morphology inside and outside water, other planets’ life forms, and robot applications outside Earth (***Rudin et al., 2022***). For simplification purposes, we choose to not explicitly control other important factors of locomotion (*i.e*., energy consumption, maneuverability) that nonlinearly interact during locomotion. In future studies, it would be important to conduct similar studies on a wider range of factors to study the shape and dynamic principles in different conditions.

Limb pattern formation - a decisive factor for locomotion - can be addressed by the study of gene expression during development (*i.e*., Hox genes implications in the number of fingers and limb evolution (***Tabin, 1992***) and limb regression in whales and snakes (***Bejder and Hall, 2002***)). Complementary, our work shows that a small to an intermediate number of modules should be a robust evolutionary outcome for organisms with directed locomotion on the ground. This result suggests the current shape’s modularity of animals (***Fischer and Blickhan, 2006; Larouche et al., 2018; Bruce, 2021***) is probably not contingent on the unique Earth’s life evolution and would consistently repeat on other planet’s life or repetitions of the tape of life on Earth (***Powell, 2020***). However, our results show that the shape’s configuration could be different from what we traditionally observe. For example, one possible configuration is to have one leg in each opposite extreme of the anterior-posterior body axis and two others positioned bilaterally symmetrical in the transverse axis (see Figure 2 - figure supplement 3). Our results point out that this limb configuration for a quadruped would be as good as the traditional one for directed locomotion on the ground with no obstacles.

The conditions that lead to the abrupt explosion of bilaterally symmetric animals and its evolutionary robustness (***Powell, 2020***) are still open questions (***Budd and Jensen, 2017; Chen et al., 2019; Heger et al., 2020***). For example, internal transport and not directed locomotion could be the original force pressuring for bilateral symmetry (***Finnerty, 2005***). Thus, the symmetry outcomes of evolution that we see today could have been different and even include the absence of symmetry. In our work, bilateral symmetry showed to be a necessary, law-like pattern in animal evolution for efficient directed locomotion purposes (***Holló, 2017***) - appearing without considering other body functions (such as the digestive system) that would bias a body to it. Beyond the gene regulatory networks or physical forces (***Holló, 2017***), the function should be a critical factor in explaining the current dominance of bilateral symmetry in animal species.

### Computationally replaying the tape of life

The biologist Stephen Jay Gould proposed a thought experiment about the contingency of natural evolution outcomes observed in Earth (***Powell, 2020***). He hypothesizes that if we could replay the tape of life, there would be very different outcomes in each repetition. Widely present biological solutions (*i.e*., two eyes) might not appear or be exceptions. The difficulty in testing the level of contingency in biological outcomes is that we do not have access to alternative evolutionary histories on Earth or other planets.

Here we propose that evolutionary physical simulations of robots can be a partial realization of Gould’s thought experiment about replaying the tape of life. This approach allows the reproduction of N>1 evolutionary process and the study of animal bodies using embodied *in silico* organisms. We expect that including other important aspects of an animal’s body as a developmental process and sensory functions could influence the shape’s outcomes with other layers of principles. Although we based our simulations on an already successful transference of *in silico* behavior to organisms made of biological tissue (***Kriegman et al., 2020***), there is an intrinsic gap between spring-mass robots modeling and animal’s bodies that is worthy of exploring to ensure the generality of our results. Other methods, such as the inclusion of rigid body elements in the simulation (possible in Voxelyze), the use of finite element modeling (FEM) (***Coevoet et al., 2019***), and the construction of physical robots (***Aguilar et al., 2016***), are important complements to this work. Beyond that, principles on other scales as in the genotypes (***Johnston et al., 2022***) and in other behavioral phenotypes (***Gomez-Marin et al., 2016***) could also be investigated. This way, with proper choices of the simulation environment, evolutionary algorithm, and encoding structures, it is possible to study the contingency level of a biological outcome given specific selection pressures. There are already evolutionary simulation studies in robots that analyze the effects of development (***Corucci et al., 2017***), environment (***Miras and Eiben, 2019; Pigozzi et al., 2023***), material properties and environment transitions (***Corucci et al., 2018***), representation (***Medvet et al., 2021***) and control (***Cheney et al., 2014***) on morphologies and behavior. Environment’s parameters as gravity - a constant variable on Earth that is difficult to test experimentally and affects life’s evolutionary outcomes (***Rayner, 2003***) - can be tested in simulation studies (***Morey-Holton, 2003***). The simulation consistency between our results with a different simulation study that also investigated the relationship of locomotion and symmetry (***C***. ***Bongard and Paul, 2000***), the behavioral transference from *in silico* to *in vivo* organisms (***Kriegman et al., 2020***), and other previous biologic studies using robots and simulations (***Miller et al., 2012; Ijspeert, 2014; Aguilar et al., 2016; Gomez-Marin and Zhang, 2022***) are evidence that *in silico* studies associated with biological questions can be used to approach Gould’s question computationally.

## Methods

The simulation experiments done in this work have three main elements: the robots (representation and material properties), their virtual environment (physic engine and simulation conditions), and the evolutionary algorithm in which we evolved them (a population optimization process). We will explain each one of them in the following sections.

The source code necessary for reproducing the computational results reported in this paper is at GitHub https://github.com/biagzi/locomotion_principles.

### Robot representation

The robots used in this work are 3D Voxel-based soft robots, which means that joined cubic deformable units (voxels) create them. Voxels are combined inside a 4x4x4 bounding box (design space) to form the robot’s body shape (Figure 1E). We did control experiments with robots within 6^3^ and 8^3^ dimensions to check for dimension size effects - and we found that the results found in 4^3^ remained valid. We choose to focus our analysis in the 4^3^ design space because we consider it the minimum coarse-grain to approach the biological question about the contingency of shape outcomes pressured for locomotion. Smaller spaces do not allow sufficient complexity in the body structures, and increasing spatial resolution reduces the extensiveness of the investigated search space.

Each robot in the simulation is represented both by genotype and phenotype representations. The phenotype consists of the robot shape (voxels configuration), control (voxels phase), and the voxels material properties. The genotype is the encoded information used to construct the pheno-type, and it was structured using different strategies - direct encoding and Compositional pattern-producing networks (CPPNs) (***Stanley, 2007***).

#### Phenotype

The phenotype is the virtual realization of the robot in the Voxelyze physics engine (***Hiller and Lipson, 2014***). Each robot’s phenotypic space of possibilities consists of *i*. voxel presence or absence in each lattice position (shape), *ii*. phase offset of each present voxel (control), and *iii*. the global stiffness, a unique stiffness value (Young’s modulus) for all voxels in the robot. Each voxel *i* has a real value *ϕ*_*i*_ *∈* [*−*1, 1] defining its phase offset in relation to the environment global signal sin 2*π*(*f t* + *ϕ*_*i*_) that controls volumetric actuation in each time *t*. The volumetric change of the oscillating voxels is ± 50% of their rest volume, and we used a fixed oscillation frequency of *f* = 2Hz (***Kriegman et al., 2020***). A fixed frequency value reduces the number of degrees of freedom in the search for solutions, but in return, it narrows the direct connection between the simulated organisms and animals. Exploring different frequency values in future work would be important to investigate the impact of varied oscillatory frequencies in the shape solutions for directed locomotion.

For each robot, the possible global stiffness values are 5e4, 5e5, 5e6 or 5e7 Pa. We based this range of values on biomechanical properties of muscle, tendons, and limbs of different aquatic and terrestrial species (***Bennett et al., 1986; He et al., 1991; Mchenry and Pell, 1995; Long, 1998; Vincent, 2012***). After the evolution, we noticed that for the water (mars and earth) environment, all the best robots had necessarily the stiffness value of 5e5 Pa (5e7 Pa). Thus, we considered these values the best stiffness for each environment. For all the analyses, we selected just the robots with the best stiffness for each gravitational environment, so we do not have stiffness variability in the results. The other material properties were kept constant during all the experiments and were based on ***Kriegman et al***. (***2020***) work that succeeded in transferring both morphology and behavior from *in silico* to *in vivo* organisms.

#### Genotype

The genotype is the encoding of the phenotype and has three independent structures - shape, control, and global stiffness. For the global stiffness of the body, we always used a Direct Encode encoding strategy that randomly chose one of the four possible values for voxel Young’s modulus. The Direct Encode genotype is a structure that directly encodes the necessary information for generating a phenotype feature. For the shape and control, we performed experiments both with the Direct Encode and the CPPN. In the cases using a Directed Encode, a matrix of size *N*^3^ (N = 4, 6, or 8) associated with each position of the *N*^3^ cartesian 3D lattice receives a random Boolean value indicating the presence or not of a voxel (shape case) or a random real value *ϕ ∈* [*−*1, 1] indicating the phase offset (control case).

The CPPNs were proposed in ***Stanley*** (***2007***) as an indirect encoding that facilitates the generation of spatial regularities. It consists of a directed network with weighted edges that multiplies the signal passing through it. The directed edges connect nodes with activation functions 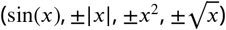 that receive the input signal and produce as output the continuous and symmetric patterns given by their functions. Thus, a fixed initial input signal passes through all the possible paths through the nodes, gets into the sigmoidal function of the output node, and gives a value in the [*−*1, 1] interval. In the case of the control genotype, this value maps directly to the phase offset values. In the case of shape genotype, a map function transforms the output to boolean values defining the presence and absence of the voxels. Using CPPN in the genotype speeds up the optimization of locomotion robots as it tends to form patterns with regularities, a feature known to be good for locomotion. However, there was the possibility that configurations that were less intuitive and regular would be as good as the commonly expected regular solutions. Doing independent simulations with CPPN and Direct Encode - a structure that does not favor regularity - allowed us to compare the results of these two search strategies.

### Virtual environment simulation

We did the simulations in the Voxelyze physics engine (***Hiller and Lipson, 2014***). It quantitatively models non-linearly deformed soft body dynamics using a mass-spring lattice. Each voxel has mass and rotational inertia values and is a lattice point with three translational and three rotational degrees of freedom. Voxels are connected by beam elements with translational and rotational stiffness, thus forming the mass-spring system. The software efficiently simulates collisions between the voxels themselves and voxels with the environment (ground or obstacles) without self-penetration.

All the simulations had the same environmental conditions except for the gravity value. The floor has no slope, and no fluidity effect was enabled. We used a Coulomb friction model with 1.0 (static) and 0.5 (kinetic) coefficients to simulate friction between the voxels and the floor. Other environment parameters were based on ***Kriegman et al***. (***2020***).

We simulated the robots during 32s, in which 2s are for settling under gravity and 30s for evaluation. We tested different time intervals for evaluation to check if it could affect the final results. When using small evaluation intervals, the robots with a faster launch that fall or trace a highly curved path usually have a higher linear displacement than robots with stable and straight trajectories. We concluded that the 30s was enough to differentiate curved and unstable from stable straight trajectories. Time intervals longer than that made no significant difference in the results and significantly delayed the experiment’s duration.

### Evolutionary algorithm

We developed an evolutionary algorithm based on ***Kriegman et al***. (***2020***). The algorithm starts generating 101 random robots that constitute the initial population. Each robot of this population has its phenotype evaluated, and 50 are selected to form the reproduction population (parents). Then, each robot of the parent group is made a copy with a mutation in the genotype, generating a new robot (offspring). The group of 50 parents, 50 offspring, and one new random individual created will constitute the new initial population of 101 robots. This process represents one generation cycle, which we repeated for 1500 generations. After testing other population sizes *P*_*s*_ and numbers of generation *G*, we found that *P*_*s*_ = 50 and *G* = 1500 are sufficient values to do an extensive search that, after a while, converges to optimal solutions. For each gravitational environment, the evolutionary process was performed 30 independent times (seeds) for the 4^3^ and 6^3^ CPPN experiments, 20 for 4^3^ Direct Encode, 10 for 6^3^ Direct Encode, and 5 for 8^3^ CPPN and Direct Encode. The different choice of seeds was due to the computational time cost and the need for an extensive search for each case. In all the cases, the number of experiments was sufficient to check our result’s robustness. Each step of the algorithm will be detailed and explained next.

#### Phenotype evaluation and selection

The phenotype evaluation consists of calculating the average speed 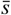 of the robot after the 30s of simulation in the Voxelyze.

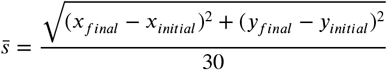

As the robots with the highest average speed are the ones that succeed in maximizing displacement and having robust dynamics (they will not tumble with time), we defined 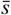 as the fitness value using it as a proxy of successful directed locomotion. Selecting for bodies that maximize speed is a common locomotion bias in natural selection, as both predators and prey and thus fecundity and mortality depend on it (***Alexander, 2006***). Other measures - such as energy efficiency - can capture distinct important aspects of the locomotion complexity (***Paez and Melo, 2014***) and would be worthy of investigating in future work.

The method used for selection was an Age-Fitness-Pareto Optimization (AFPO) (***Schmidt and Lipson, 2011***). AFPO is a multi-objective method that uses fitness and age as a criterion to select the robots that will survive and constitute the reproduction population. The new randomly created robots start at age 0. The offspring created by mutating a robot of the reproduction population inherit the parent’s age. After generating the offspring but before adding a new random robot to the population, we increment the age of all the 100 robots by one. Thus, age measures how much time the algorithm passes searching a specific region of the space of possibilities. The robots are classified in dominance levels using the Pareto fronts of multi-objective optimization, maximizing fitness, and minimizing age.

#### Genotype mutation

In each generation, the offsprings are a mutated copy of each robot of the reproduction population. To increase the impact of mutations on shape variability inside the population, we used a diversity filter that actuates with a probability of 1/2 in each generation. If applied, the filter ensures that the offspring group in a generation will not contain robots with shape similarity ≥ 95%. If the diversity filter is not applied, the algorithm includes the first offspring of each parent without any restrictions. This way, mutations that have both detailed (for body optimization) and large (for keeping shape exploration) effects can happen. In each parent mutation, the shape, control, and global stiffness genotype structures have a 1/2, 1/2, and 1/5 chance of being selected for mutation. Thus, the type of mutations can differ in each new offspring.

The mutation in the global stiffness genotype (Direct Encode) is simply a random choice of one of the four options (5e4, 5e5, 5e6, or 5e7 Pa) as long as it is not equal to the previous stiffness. The mutation in shape and control as Direct Encode is a change of values in places of the lattice randomly chosen. In the case of shape and control as CPPN, it happens by adding, modifying, or removing a randomly chosen node or edge of the network. Each of the six kinds of network mutation (add node/edge, modify node/edge, or remove node/edge) has a 1/6 probability of being selected (if none are selected, one of them is applied). If the phenotype has not changed after a network mutation, another mutation happens until 1500 attempts. If the 1500 attempts fail to change the phenotype, all 1500 neutral mutations are applied.

### Environment Transference

After running experiments with the evolutionary algorithm in different environments, we did the environment transference of the best robots of size 4^3^ and CPPN genotype (Figure 4B). First, we sorted all robots generated in all 30 seeds by their fitness. Then, we select the 50 best robots with different shapes for each environment. Finally, we considered different shapes with at least four different voxels when compared with each other. The comparison consisted in counting the number of different voxels of each two shapes when one of them is reflected and rotated in all its possibilities. The possibilities are *i*. the original position and its four rotations in the z-axis (0, 90, 180, 270°), *ii*. the reflection in the x-z plane and its four rotations in the z-axis, and *iii*. the reflection in the y-z plane and its for rotations in the z-axis (4+4+4). Thus, the shapes are different if the closest comparison (minimum value of 12 possibilities) is greater than 4.

For each original gravitational environment, we transferred all the 50 best shapes to the other two gravities and their original one (control trial). The transference consisted of optimizing each shape’s phase offset (control of the body) in the new gravitational environment. The optimization consisted of evolving the robot, with fixed shape and stiffness, in the evolutionary algorithm described above. The optimization used initial populations of 10 robots with five new random controls added in each cycle and took 200 generations. We checked that this generation number was sufficient to find the best way to move the body (there is a fitness converge). We did not use the diversity alternation filter, as its function is to increase shape diversity, and in this optimization we fixed the shapes. We used a fixed voxel stiffness value of 5e5 Pa for water and 5e7 Pa for mars and earth. After the 200 generations, we choose the best control (phase offset solution) for analyzing the performance in the new environment.

### Modularity measurement by clustering

We used an algorithm to cluster the neighboring voxels with similar phase offset values inside each robot’s body, and this was our measure of modularity. We used DBSCAN (***Pedregosa, F***. ***et al., 2011***) as it uses parameters (minsample, eps) that have a meaning in our system and do not need to be blindly guessed. DBSCAN uses distances between the nearest points to define the density regions in the sample space. Each point of our sample is a four-dimensional array that represents the position and phase information (*x*_*i*_, *y*_*i*_, *z*_*i*_, *ϕ*_*i*_) of each voxel *i*. After several tests, we developed an algorithm with three clustering steps and a choice of parameters that kept the clustered robot performance similar to the originals by 90% on average.

The first step of the algorithm clusters just the phase offset. It uses *minsample* = 2 (two voxels are sufficient to be considered a cluster) and *eps* = 0.2 (10% of the total range, [*−*1, 1], of phase offset values). After the first clustering, the algorithm checks if there is no cluster with a phase offset difference larger than 0.2 inside its members. If there is not, it follows to the next step. If there is, it stays in a coarse grain clustering that repeatedly uses *eps* = *eps −* 1*/*100 until the cluster results pass the check. It ensures that clusters with very different phase offsets inside it will not occur without the risk of wrongly fragmenting too much of the clustering.

After the first step, the voxels receive their cluster’s average phase offset value *⟨ϕ*_*i*_*⟩*. The second step has a similar structure of the first one, but it clusters position and mean phase offset using the sample of four dimension arrays (*x*_*i*_, *y*_*i*_, *z*_*i*_, 10 × *⟨ϕ*_*i*_*⟩*) and initial *eps* = 1.2. The factor 10 brings the *⟨ϕ*_*i*_*⟩* to the same order of magnitude of position, as *eps* = 1.2 apply equally to the four dimensions. This step uses *eps* = *eps −*1*/*10 to do the coarse grain clustering if the result does not pass the check (the same as in step 1). After the second step, the third step checks for outliers (noisy samples). If there is, each one is independently considered a cluster of 1 voxel or grouped in one of their direct neighbor clusters if they have a phase offset similar enough *≤* 0.3.

We constructed the robot’s modular representation using the cluster’s members’ average phase offset and position. The modules are linked if they have voxel members directly connected.

#### Module connectivity

Let *M* be the number of modules (nodes) of the robot and *c*_*i*_ the number of connections (edges) of each module *i* with *i* = 1, …, *M*. Then, we defined the average degree as:

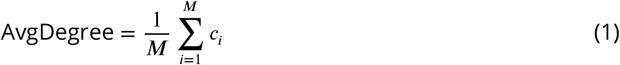

#### Number of modules amplitude

In all the plots showed in Figure 2D and Figure 2 - figure supplement 1 the fitness layers are normalized together. Thus, we can observe that the worst fitness layers (40-20% and 20-0%) constitute a larger sample of robots than the best fitness layers (100-80% and 80-60%) in all the experiments and all the environments. This difference in sample sizes is expected, as the higher fitness results from the specialization of a small subset of survivor robots given a large number of bad trials that are extinct. In the specific case of the number of modules analysis, the layers distributions have a similar shape except for a tail present in a small density. Thus, we checked if the amplitude difference between the layers was significant or was due to extremely rare outliers present in the worst layers because of their bigger sample sizes. For that, we did a bootstrap of the sample for each environment and each experiment, with N = 10000. In each one of the random sampling with replacement, we saved the number of modules amplitude for each layer. This way, after the boot-strapping we had a distribution of the number of modules amplitude (Figure 2 - figure supplement 2).

---

**Figure 2–Figure supplement 1.**
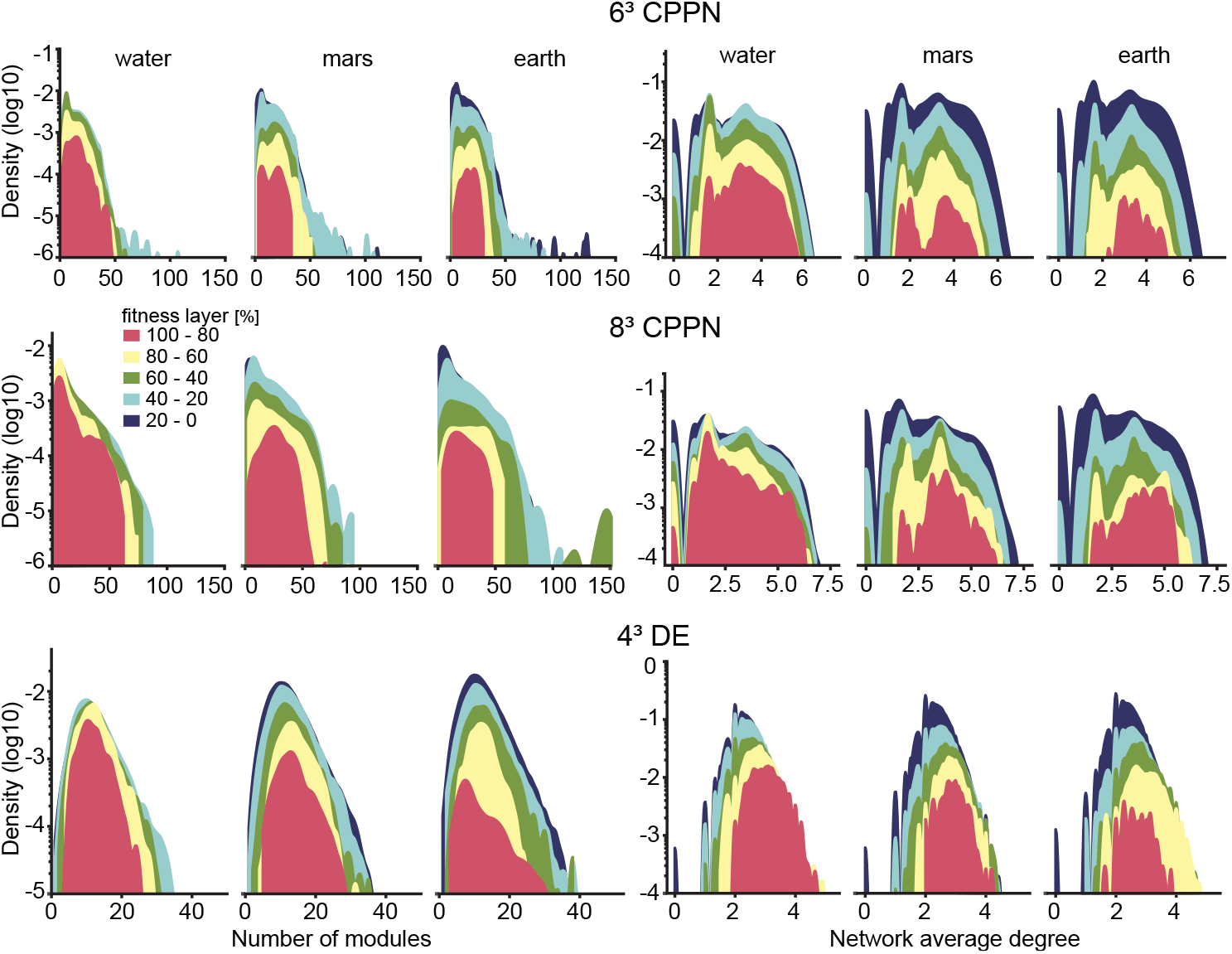
The results of the three experiments (6^3^ CPPN, 8^3^ CPPN, 4^3^ DE) confirm the results of 4^3^ CPPN that the best robots (100-80% fitness layer in red) typically have an intermediate number of modules and average degree compared to the other layers. Thus, the principles found seem to hold for different genotypes and sizes of robots.

**Figure 2–Figure supplement 2.**
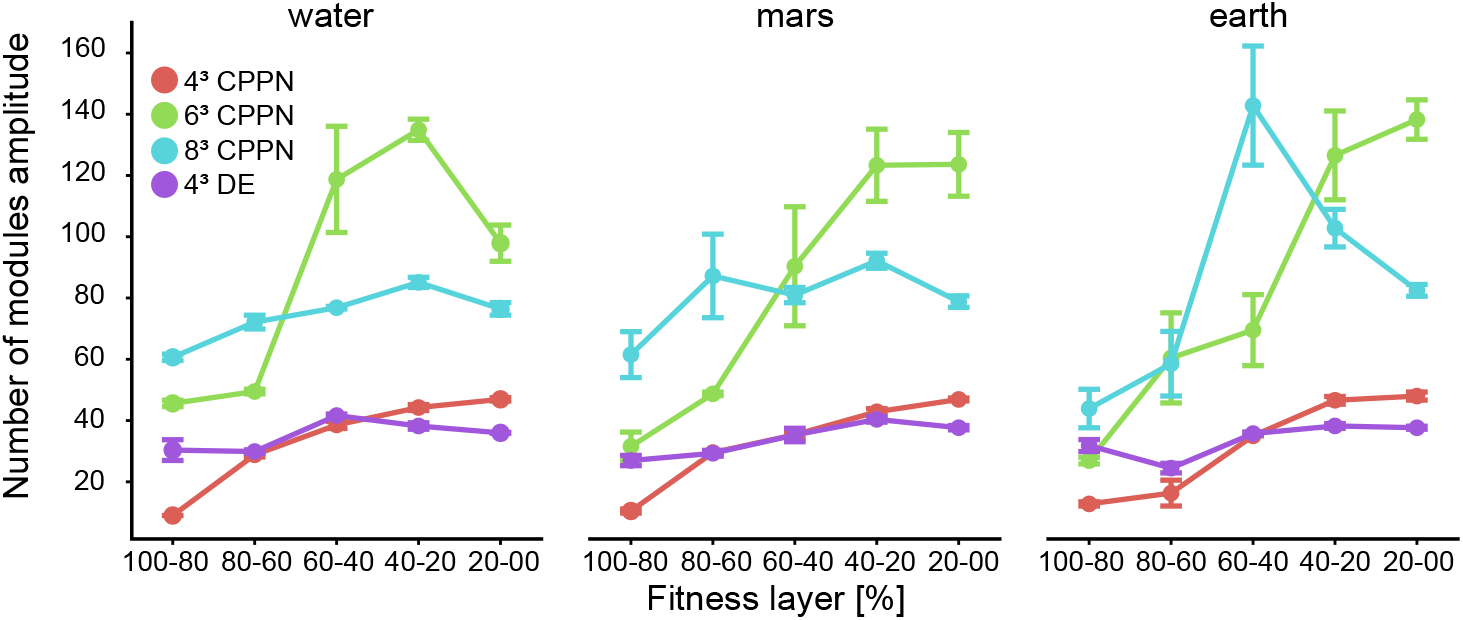
The results of the three experiments (6^3^ CPPN, 8^3^ CPPN, 4^3^ DE) confirm the results of 4^3^ CPPN that the best robots (100-80% and 80-60% fitness layers) restricts to a small amplitude of values compared to the worst layers (40-20% and 20-0%). We did a bootstrap of the sample for each environment and each experiment, with N = 10000. We plotted the difference of the extreme values (amplitude) for each layer distribution. The points are the mean value of the distribution of amplitude values after N=10000 bootstrapping.

**Figure 2–Figure supplement 3.**
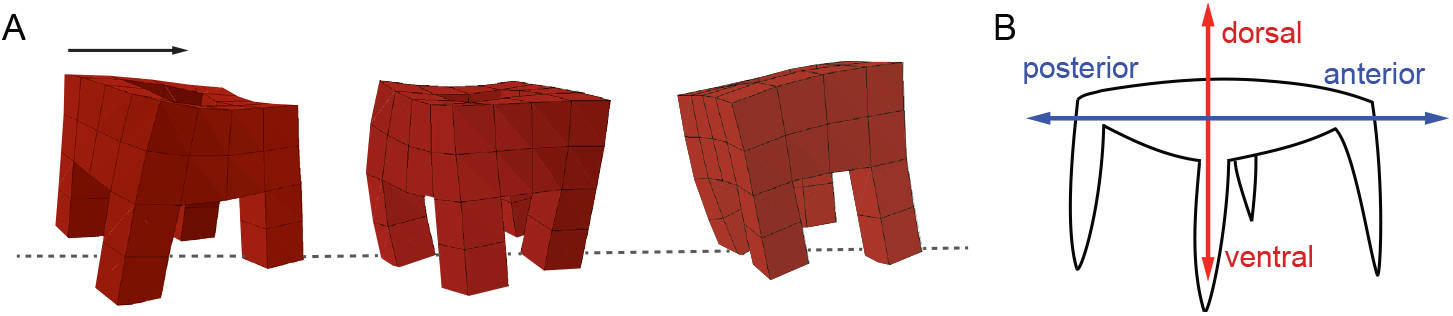
Alternative limbs configuration for bilateral quadrupeds.

**Figure 3–Figure supplement 1.**
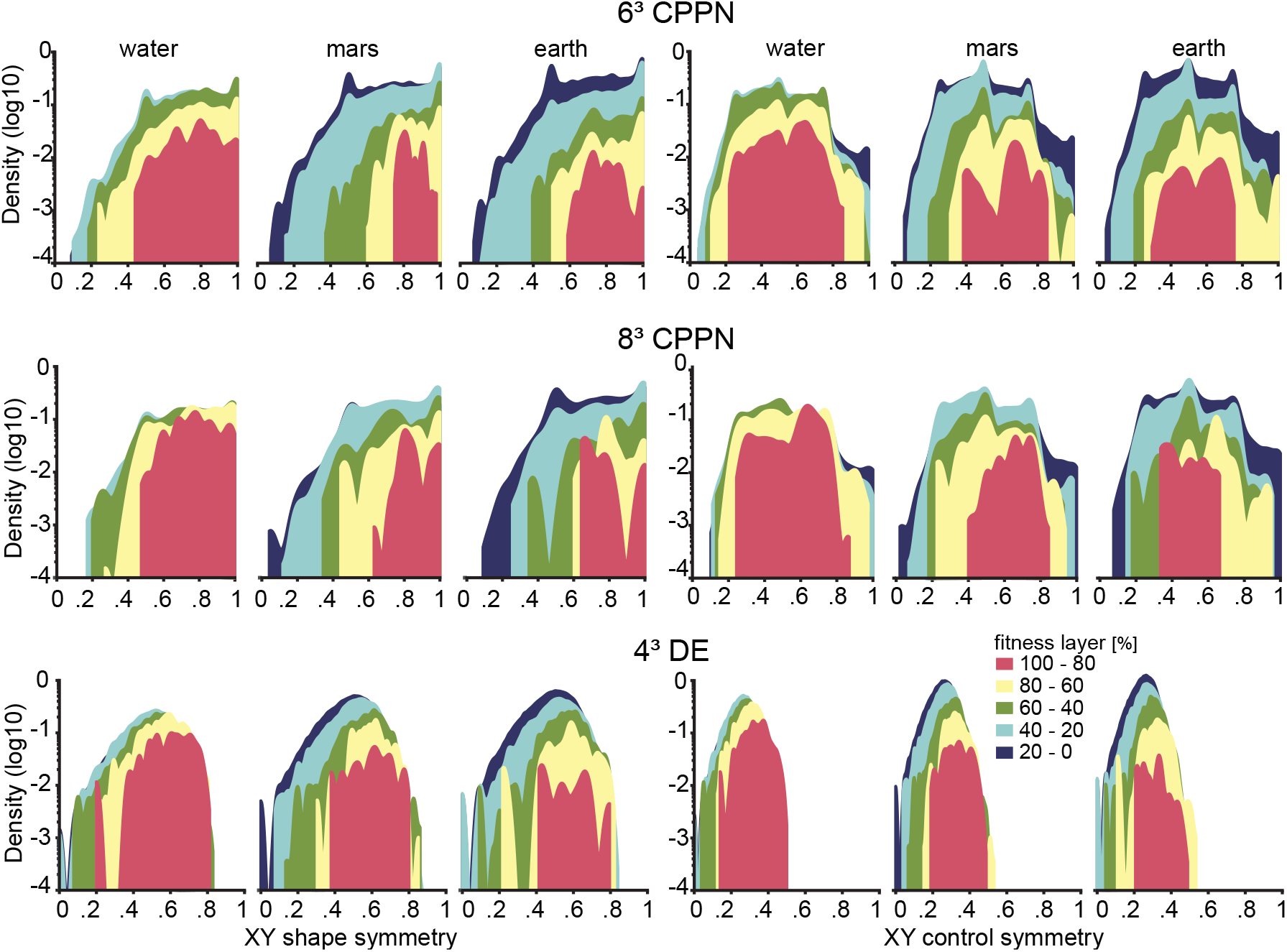
The three experiments (6^3^ CPPN, 8^3^ CPPN, 4^3^ DE) confirm the results of 4^3^ CPPN that the best robots (100-80% fitness layer in red) typically are shifted to higher shape and control symmetry values compared to the other layers. Besides, in all the cases, there is a break of total symmetry in the control symmetry. Thus, the principles found are true even when using different genotypes and robot sizes.

## Notes

### Competing Interest Statement

The authors have declared no competing interest.

### Summary of Updates

The Results, Discussion, and Methods sections were updated to make the text clearer and more precise concerning the metrics and definitions used, the hypothesis, and the extent of the conclusions.

